# Nationally endorsed learning objectives to improve course design in introductory biology

**DOI:** 10.1101/2023.10.25.563732

**Authors:** Kelly M Hennessey, Scott Freeman

**Author notes:** Corresponding author (KMH). These authors contributed equally to this work.

## Abstract

Introductory biology for majors is one of the most consequential courses in STEM, with annual enrollments of several hundred thousand students in the United States alone. To support increased student success and meet current and projected needs for qualified STEM professionals, it will be crucial to redesign majors biology by using explicit learning objectives (LOs) that can be aligned with assessments and active learning exercises. When a course is designed in this way, students have opportunities for the practice and support they need to learn, and instructors can collect the evidence they need to evaluate whether students have mastered key concepts and skills. Following an iterative process of review, revision, and evaluation, which included input from over 800 biology instructors around the country, we produced a nationally endorsed set of lesson-level LOs for a year-long introductory biology for major’s course. These LOs are granular enough to support individual class sessions and provide instructors with a framework for course design that is directly connected to the broad themes in *Vision and Change* and the general statements in the BioCore and BioSkills Guides. Instructors can implement backward course design by aligning these community endorsed LOs with daily and weekly learning activities and with formative and summative assessments.

## Introduction

Research has shown that active learning increases achievement for all students [1] and especially for populations underrepresented in STEM [2]. An important next frontier for continued improvement in student performance—a requirement for the U.S. to meet current and projected needs for qualified STEM professionals [3]—is to align improvements in active learning exercises and other evidence-based practices with specific learning objectives (LOs), then measure student progress on those LOs with aligned assessments. Work in cognitive sciences comparing the methods of novices and experts in a field has identified a common trait among novice learners: They are not yet capable of evaluating the relative importance of information, meaning that they cannot distinguish less-important from more-important information [4].

Because they are new to the field, novices lack the experience in problem solving and background in key organizing concepts and frameworks that experts rely on to pick out key information efficiently [4]. Theory suggests that LOs provided by experts will help novices with this task, as they are explicit statements about the knowledge and skills that are important to master [5]. By employing the Backward Design framework, instructors begin with clear, specific learning objectives and then work backward to create instructional activities and assessments that are closely aligned with these objectives [6]. This alignment helps students develop the understanding and skills necessary to enhance their performance. When learning objectives and assessments are aligned, students can focus their studying, benefiting from deliberate practice and becoming more efficient and effective [6]. This approach ensures that assessments accurately evaluate whether students have achieved the learning objectives. Both elements of the theory predict that careful use of well-designed LOs will increase opportunities for equity and improve student outcomes [7].

The rapid pace of scientific discoveries has broadened the field of biology, presenting a challenge for biology instructors in deciding what to include in a year-long introductory college biology course [8]. However, biology educators have made important progress toward the goal of developing a cohesive set of LOs. The *Vision and Change* initiative [9], for example, was transformative in life sciences education because it provided a set of overall concepts and competencies and themes that broke with the “cover all the content” approaches that dominated instruction previously. Because it was developed with input from over 500 biology faculty, *Vision and Change* was presented as a broad national consensus.

The effort to develop comprehensive course design criteria that began with *Vision and Change* [9] continued with the publication of the BioCore Guide [8] and the BioSkills Guide [10]. Each of these guides added another level of sophistication to the *Vision and Change* framework by providing statements that elaborated on the general goals laid out in the *Vision and Change* report. In the BioCore Guide, for example, *Vision and Change*’s unifying concept of Evolution is broken down into focused statements such as “Most organisms have anatomical and physiological traits that tend to increase their fitness for a particular environment.” Both guides were developed with input from hundreds of biology instructors and thus—like *Vision and Change*—represent a broad national consensus.

The publication of the BioCore Guide and BioSkills Guide inspired the development and testing of a series of programmatic-level assessments. These Bio-MAPS instruments are given to students at different points in their undergraduate careers, culminating at graduation, to evaluate the effectiveness of the overall curriculum at particular institutions [11–14].

Thoughtful, well-trained instructors anchor their courses with these types of broad-based, term-long learning outcomes that unify topics in the curriculum. But high- and mid-level course or programmatic goals, like those encapsulated in the *Vision and Change* report and BioCore Guide and BioSkills Guide, lack the granularity needed for daily or weekly learning objectives (Fig 1). Lesson-level LOs complement programmatic- or course-level learning outcomes by stating what instructors expect students to know and be able to do by the end of a specific class session [5]. As a result, they are granular enough to be aligned with active learning exercises and individual assessment items.

**Fig 1.**
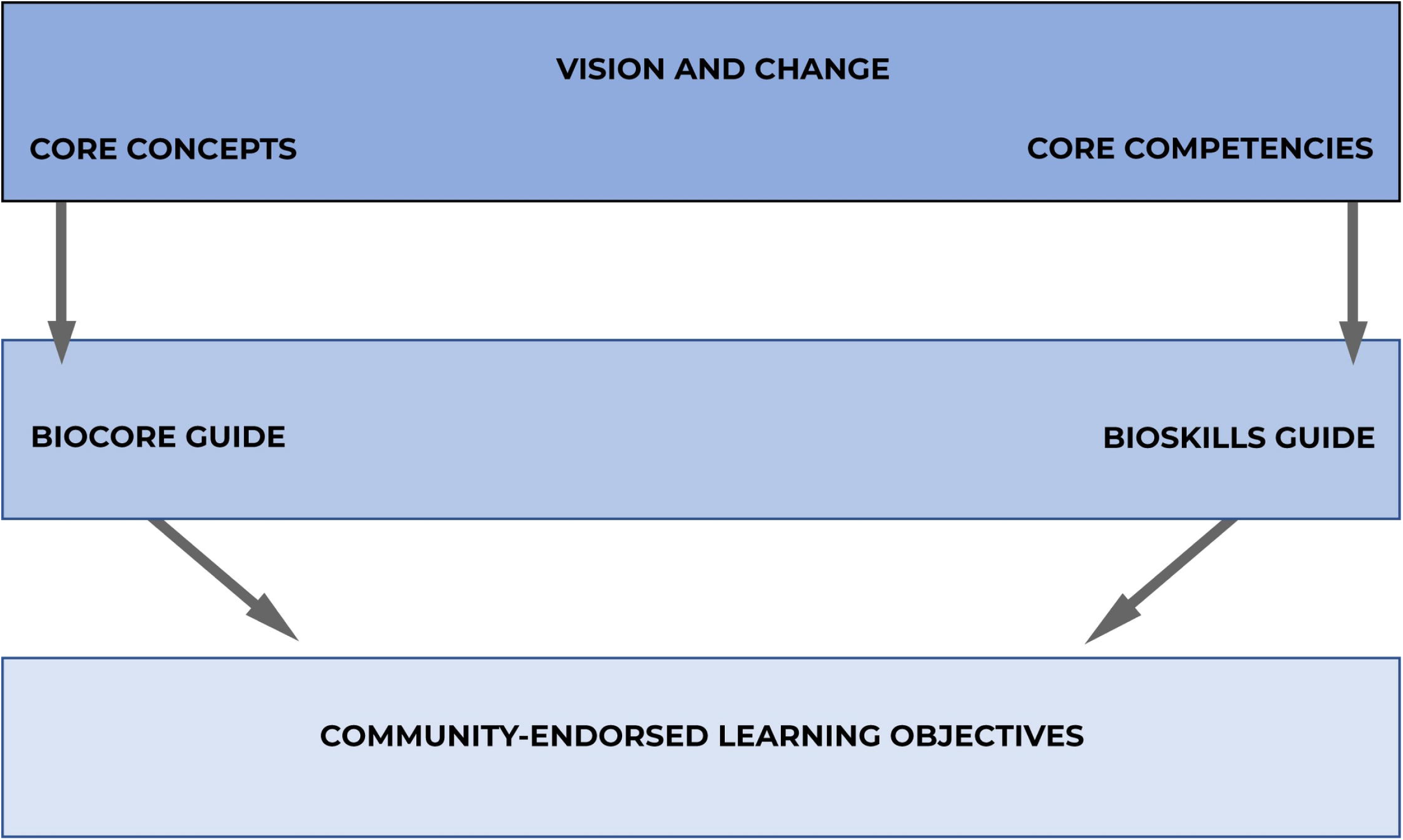
Hierarchy of life-science course goals and objectives.

### The role of learning objectives

Many terms in the literature are used interchangeably to describe learning expectations. Examples such as “learning outcomes,” “learning goals,” “course outcomes,” and “course goals” convey similar meaning but lead to confusion for students and instructors alike. To provide consistency and clarity, we follow Orr et al. (2022) by referring to learning objectives, abbreviated as LOs, as statements of what students should know and be able to do at the end of a specific class session and “outcomes” as results that are measured at the end of a unit, course, or program.

Effective curricular design begins by identifying LOs as the desired end result [6]. This backward design approach encourages faculty to start with clear statements of overall learning goals for a course that are aligned with lesson-level LOs. Ideally, this step would involve a combination of nationally endorsed standards and instructor-written aims that customize the course to the institution and student population. Based on statements of what students should know and be able to do, faculty can then design teaching activities that will support students as they achieve mastery. In addition, faculty can write questions and exercises for formative and summative assessments that align with their LOs and classroom practice. We envision dynamic and reciprocal integrated course design that emphasizes the relationships among LOs, teaching practice, and assessment.

Clear and thoughtful LOs help students better understand lesson and course expectations and the projected directions for their learning [5,15]. In a well-designed science course, students and instructors make explicit use of these LOs, use in-class activities and out-of-class assignments to master them, and evaluate progress with carefully constructed formative and summative assessments [16]. To maximize student progress, all three course components—the LOs, the learning activities, and the assessments—should be aligned and mutually reinforcing.

In addition, mastering a set of LOs provides a foundation for vertical alignment among courses in a major. This cascade effect means that a failure to meet LOs at the introductory level can impact a student’s ability to understand more challenging concepts later in their undergraduate careers. For example, if students do not master an introductory-level LO on how meiosis generates genetic variation in gametes and offspring, they may not be able to address an LO on evaluating hypotheses for the evolution of sex in an upper-level Genetics or Evolution course.

In all-too-many cases, however, instructors design their courses in the complete absence of LOs and organize instruction around content coverage—often simply following textbook treatments [17]. When this happens, upper-division instructors and graduate and professional schools may inadvertently re-teach material unnecessarily, as they cannot be confident about what students have already been required to master. When instructors do use LOs—often because they have an administrative requirement to either develop their own or to use learning objectives that are mandated—they can vary in quality and in their level of cognitive challenge. As a result, learning objectives may vary enough across institutions that courses do not support articulation agreements, complicating the ability of transfer students to integrate into a new institution.

### Aligning course outcomes with assessments

Research has documented a glaring and persistent mismatch between course-level learning goals and the assessment items actually used in courses, especially exam questions used to assign grades. Even instructors who have been trained in evidence-based instruction [18] and who embrace the unifying conceptual frameworks outlined in *Vision and Change* [8, 9] and the competencies-based focus of Training Future Physicians [19] routinely give exams that predominantly assess recall or low-level conceptual understanding [20–23]. Assessments that primarily reward rote memorization do not cultivate competence in the analytical and lifelong learning skills needed to thrive in a 21^st^-century economy [3].

Programmatic assessments, such as the Bio-MAPS assessments [11], which were developed to assess the effectiveness of a general biology curriculum in teaching the five core concepts outlined by *Vision and Change*, are an important step in designing a four-year curriculum.

However, programmatic assessments such as these are too broad to serve as lesson-level assessments. Developing lesson-level LOs and assessment items that align with broader course LOs remains a major challenge for the community [17, 24].

### Aligning lesson level LOs with activities and assessments

Introductory Biology for Majors is one of the most consequential courses in undergraduate STEM education, with annual enrollments of several hundred thousand students per year in the United States alone [25]. To support instructors who want to implement backward design in their course sequence, we facilitated the development of a set of lesson-level LOs that were then evaluated by over 700 instructors from a wide array of institution types. These LOs are designed to be specific enough to guide the design of individual teaching activities and out-of-class assignments and align with individual formative and summative assessment items. By supporting close alignment between LOs, teaching practice, and assessment items, the set of community-endorsed, lesson-level LOs for introductory biology published here completes the course design hierarchy (Fig 1) and supports instructors who want to implement backward course design by aligning LOs with instructional strategies and assessments.

## Materials and methods

This study was designed to support our research goal of developing LOs that align with the broader course and programmatic outcomes articulated in *Vision and Change* [9]. We also intended for the LOs to support *Vision and Change’s* general goals of reducing content coverage and rote memorization in introductory biology for majors and placing increased emphasis on higher-order cognitive skills and acquisition of professional competencies—including those articulated in the BioSkills Guide [9, 10]. We followed Orr et al., (2022) in terms of best practices in writing LOs, and specifically aimed to make the LOs granular enough that instructors could use them to design a single class session and directed enough that students would be able to write self-test items to support independent study. We also followed established practice in seeking to establish endorsed LOs intended to comprise 75% of an introductory course for majors, leaving 25% for instructors to develop on their own (e.g., [26]). The 75:25 split is designed to provide instructors with latitude to customize course content for their program and student population, so that courses have a common core of nationally endorsed LOs along with a significant percentage of LOs customized for the local context.

The methods we pursued were intended to mimic, as closely as possible, the approach developed by the authors of the *Vision and Change* report, the BioCore Guide, the BioSkills Guide, and the ASM Curriculum Guidelines [8–10, 27]. Specifically, we sought to pair an extensive and iterative development phase that engaged both teaching practitioners and education researchers, with a large national evaluation effort designed to identify a broad national endorsement for core LOs that are essential for all introductory biology courses for majors.

We received written notification from the University of Washington Human Subjects Division stating that this proposed activity is human subjects research that qualifies for exempt status (IRB ID: STUDY00010583). Therefore, this research is exempt from the federal human subjects regulations, including the requirement for IRB approval and continuing review.

### Overview of process

This project was divided into two broad phases: a development phase and an evaluation phase (Fig 2). During the development phase, candidate LOs went through multiple rounds of evaluation and revision. In total, there were six different groups of researcher-instructors that participated in the development phase. Each group involved different teams of evaluators, all of whom shared instructional expertise in life sciences content and experience with biology education research.

- Group One was composed of 11 education research scientists who regularly advise the BioInteractive program at the Howard Hughes Medical Institute (HHMI) on curriculum development.
- Group Two consisted of four biology researcher-educators who were recruited because of their extensive background in writing and evaluating LOs.
- Group Three included four experts in survey design, assessment design, and course redesign.
- Group Four was made up of two experts with extensive experience in writing LOs and in assessment design.
- Group Five comprised 20 discipline-based education researchers who had published extensively on LOs, teaching practice, and course transformation efforts.
- Group Six was comprised of this manuscript’s authors (KH, SF).

**Fig 2.**
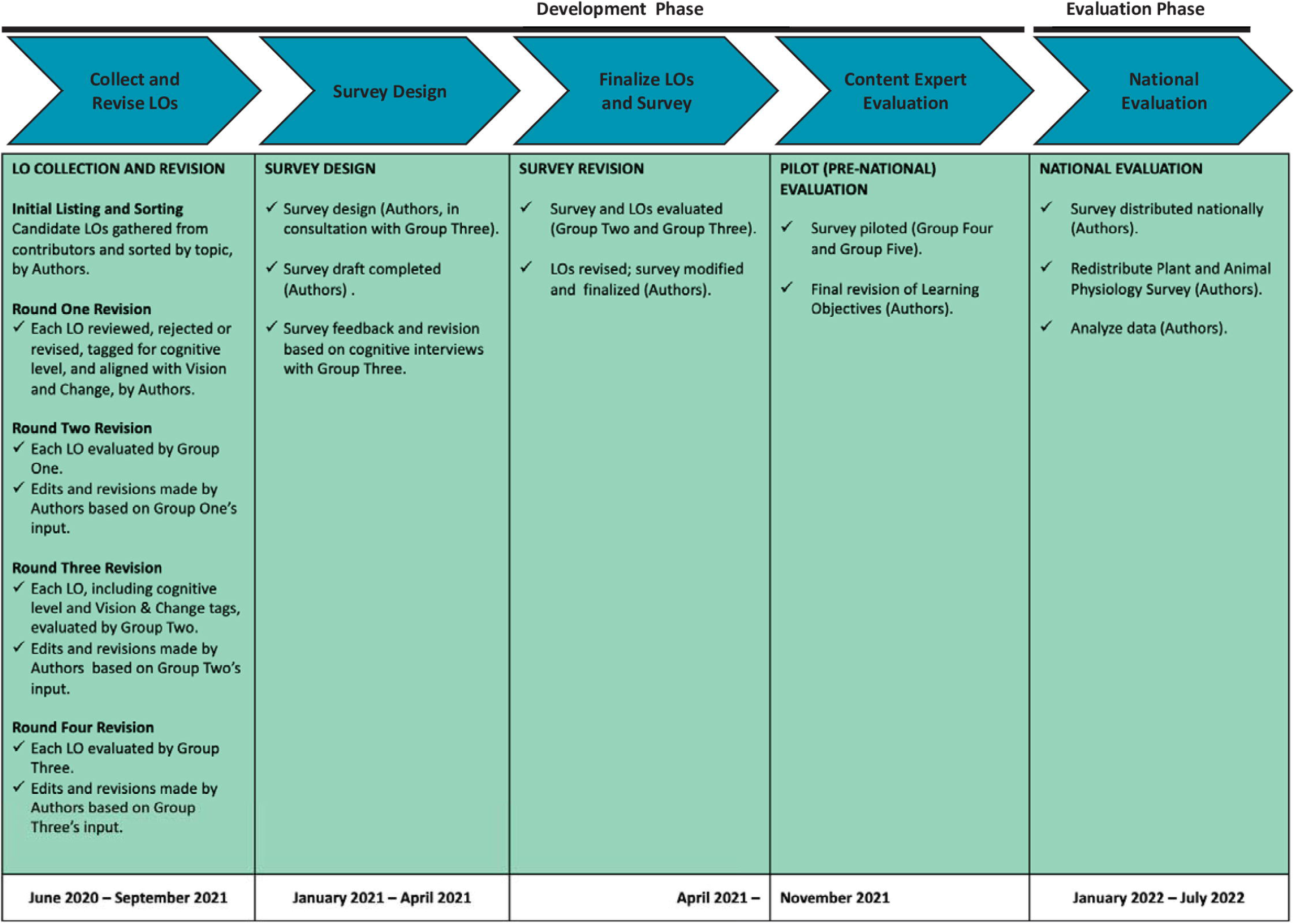
Project overview: developing learning objectives for introductory biology for majors.

For clarity, the authors will be referred to as “we” or “authors” and the remaining groups referred to by number (Group One, Group Two, etc.)

Once the development phase was complete, the LOs went through an evaluation process based on distributing a national survey to a convenience sample of instructors who teach introductory biology for majors. These faculty assessed and rated each LO as essential or nonessential for introductory biology for majors’ courses, while recognizing that each faculty member would also contribute their own LOs to customize their course to their institution and student population.

### Development Phase

The development phase started with a list of over 3000 LOs for introductory major’s biology that had been solicited by colleagues at HHMI’s Science Education program, using their listserv of BioInteractive curriculum developers and users. As Fig 3 shows, the 63 faculty who sent LOs in response to this request from HHMI represented an array of institution types.

**Fig 3.**
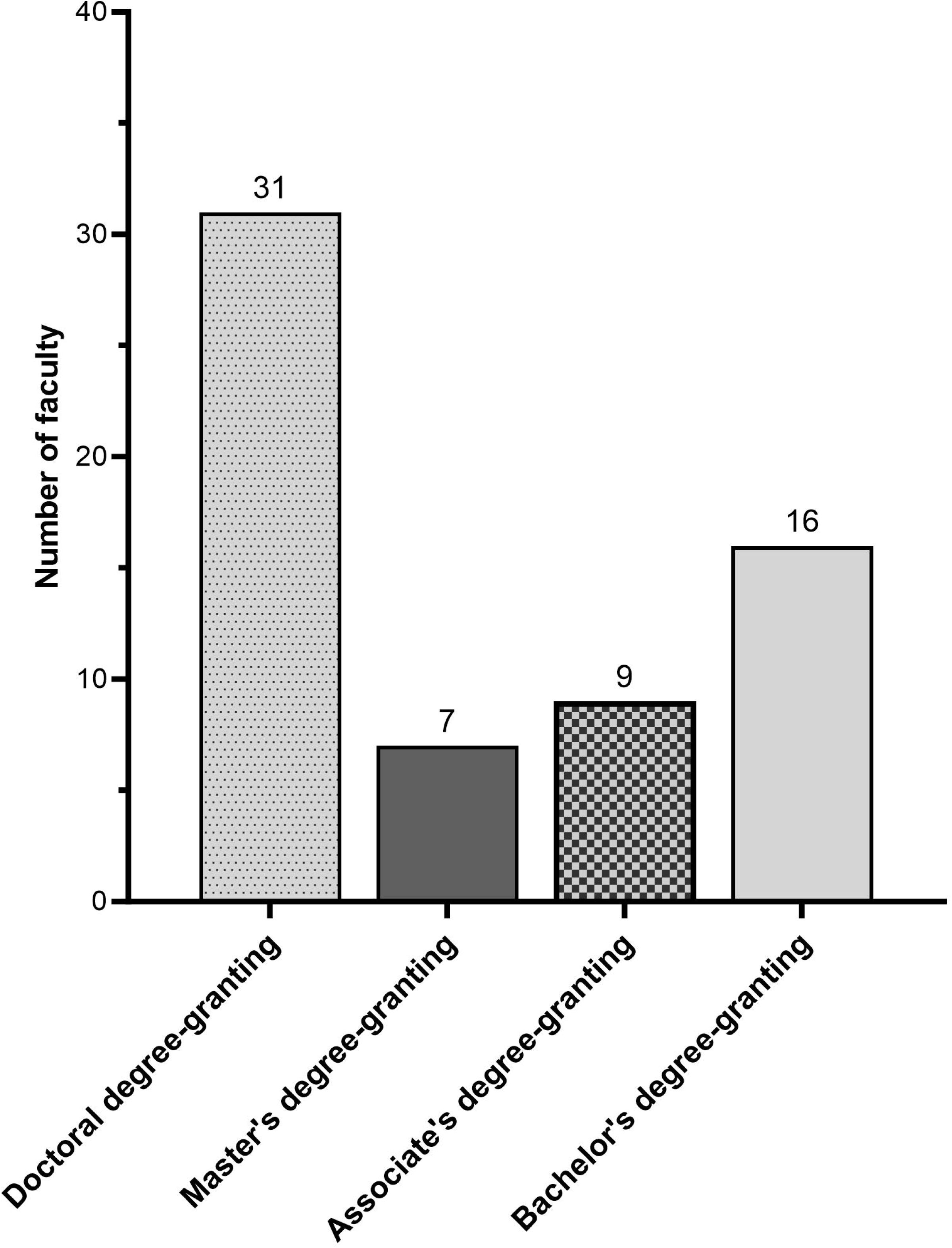
Institutional affiliations of instructors who submitted candidate LOs.

We sorted this initial list of candidate LOs into seven units based on the organization and content of several college biology textbooks including two major introductory biology textbooks [28,29]. The units were Biochemistry of Life, Cells, Genetics, Evolution, Biodiversity of Life, Plant and Animal Physiology, and Ecology. The LOs in each unit were then subdivided by topics that corresponded roughly to sections and subsections in these texts.

Once the initial list was organized by topic, we combined redundant statements and removed entries that were either beyond the scope of introductory biology or unrelated to biology content, as judged by the material covered in leading textbooks [28,29] for the course. This filtering process reduced the list of over 3000 candidate learning objectives to about 1200.

Our initial round of screening and organizing candidate LOs continued with a revision step. The need for this work arose when an initial evaluation of the draft LOs raised an important concern: very few LOs from the initial submissions asked students to apply, analyze, evaluate, or synthesize concepts. Stated another way, practices that are considered higher-order cognitive skills (HOCS) on Bloom’s taxonomy of learning were greatly underrepresented in the initial set of LOs. To address this situation, we re-wrote many LOs that focused on the lower-order cognitive skills (LOCS) of recalling vocabulary and understanding concepts with the goal of asking students to apply, analyze, evaluate, or synthesize the stated ideas and skills. After this step, each author independently rated each candidate LO as representing either LOCS or HOCS, paired LOCS and HOCS LOs on the same concept whenever possible, and aligned each LO to one or more of the concepts and competencies articulated in the final *Vision and Change* Report [9] as well as one or more statements in the BioCore Guide and BioSkills Guide [8,10]. Finally, the authors met to discuss and reach agreement in each LOCS versus HOCS designation, proposed LOCS-HOCS pairing, and *Vision and Change*, BioCore Guide, and BioSkills Guide tags.

The candidate LOs that emerged from this initial work were sent to Group One for revision as noted in Fig 2. Each of the experts in Group One reviewed the wording of every candidate LO, made suggestions for revision, and evaluated each as essential or non-essential for introductory biology for major’s students. Once this step was complete, we revised the learning objectives based on Group One’s feedback.

Additional rounds of critique and revision were completed by the four experts in Group Two, the four experts in Group Three, and the authors using the same process (Fig 2). In total, each LO went through four rounds of revision during the study’s “LO Collection and Revision” step (Fig 2).

To lay the groundwork for a large national survey focused on evaluating whether or not each LO was considered essential for introductory biology, we followed best practices in survey design [30] and the principles of social design theory during the Survey Design step highlighted in Fig 2. After developing a preliminary design for the survey in the Qualtrics platform, we engaged Group Three in providing feedback on survey design via written comments and cognitive interviews. We revised the general survey format based on these recommendations and in response to information from think-aloud interviews we conducted with four other colleagues as they took the initial version of the survey. In addition, to minimize cognitive load on respondents and maximize survey responses during the national evaluation step, units were separated into blocks of 18-24 LOs. We added this step based on preliminary data indicating that evaluating a block this size would take an instructor an average of 15 minutes to complete, and that limiting effort to 15 minutes would maximize the quality of responses by minimizing survey fatigue.

To further develop the survey, we had each member of Group Two and Group Three take at least one block of the draft survey and comment on both the survey design and the LOs. We revised both the survey and the LOs based on this feedback, resulting in a final format and structure for the survey instrument.

As a final check prior to the large-scale national survey, the pilot survey was distributed to Groups Four and Five with a request for detailed written feedback on both the survey questions and the LOs themselves. The survey and learning objectives were finalized by the authors based on responses gathered during this pilot survey. This “Content Expert Evaluation” step (see Fig 2) closed out the project’s Development phase and resulted in a total of 352 candidate LOs. In total, each of these candidate LOs was assessed and revised an average of 13 times over the course of this work.

### Evaluation Phase

We invited over 13,000 biology instructors to participate in the national evaluation step. We solicited participation through direct emails and the BioInteractive listserv and encouraged recipients to share the invitation widely. A complete copy of the survey is available in the Supporting Information.

Our effort to seek responses from as broad a spectrum of the teaching community as possible was inspired by the methods pioneered in the BioCore Guide [8] and has been variously described as grassroots, bottom-up, or faculty-first. Research has shown that community-based approaches like this are more effective at promoting buy-in and adoption of a research product than traditional Delphi strategies, which rely on top-down mandates from a small group of experts [31].

To participate in the survey, respondents were first asked to confirm that they had taught an introductory biology course for majors. Once respondents entered the survey itself, they self-selected a unit to evaluate based on their current or prior teaching experience and expertise. We had broken each unit into 1-3 blocks of LOs, based on the total number assigned to that unit, and participants received one block at random. Respondents then evaluated each LO as “essential” or “not essential” to the course they teach. They were also invited to share any feedback about the content or wording of the LOs they evaluated. Respondents were then given an opportunity to return to the survey start and evaluate an additional block of LOs. Before closing out, the survey also collected institutional and demographic data from each respondent. But to support evaluators who wished to critique the LOs, all surveys were completed anonymously.

It is important to recognize that due to survey formatting constraints, respondents were able to view and evaluate only one block of LOs at a time. As a result, they could not see all of the LOs proposed for an entire unit—a step that we hypothesized would support our goal of gathering data on LOs as “stand-alone” teaching goals. In addition, respondents evaluated each LO as a single item at a time on their computer screens, meaning that they were unaware of the pairing between LOCS and HOCS LOs. Finally, respondents did not see any of the data on alignment with Bloom’s level, *Vision and Change* Core Concepts and Competencies, BioCore Guide statements, or BioSkills Guide statements.

After our initial solicitation of participants and a preliminary assessment of the completed surveys, we realized that the number of evaluators for the Physiology unit was low. In response, we made a directed appeal to instructors with teaching expertise in animal or plant physiology who were on the BioInteractive listserv and had not already participated in the survey to complete those survey blocks. After including the data from these follow-on respondents, the total number of survey participants exceeded 700. Although the number of evaluators to rate each LO varied widely—from 25-65—the average number of raters for each LO was 38.

## Results

Using a national survey, we collected evidence on whether the teaching community considered each of 352 candidate LOs as essential in a one-year introductory biology sequence for majors. Of the 706 evaluators who participated in the survey, 609 provided data on their institution and demography (Fig 4a), with 33% of respondents coming from Associate’s degree-granting institutions, 28% from Bachelor’s degree-granting institutions, 14% from Master’s degree-granting institutions, and 25% from Doctoral degree-granting institutions. Although we did not ask whether respondents had full-time or part-time appointments, roughly 64% of respondents stated that they were assistant, associate, or full professors while 35% identified themselves as lecturers, instructors, or teaching faculty (Fig 4b). Almost half of the survey participants (46%) had no DBER experience (Fig 4c) and almost 57% were female (Fig 4d). Slightly over 77% of respondents were white (Fig 4e).

**Fig 4.**
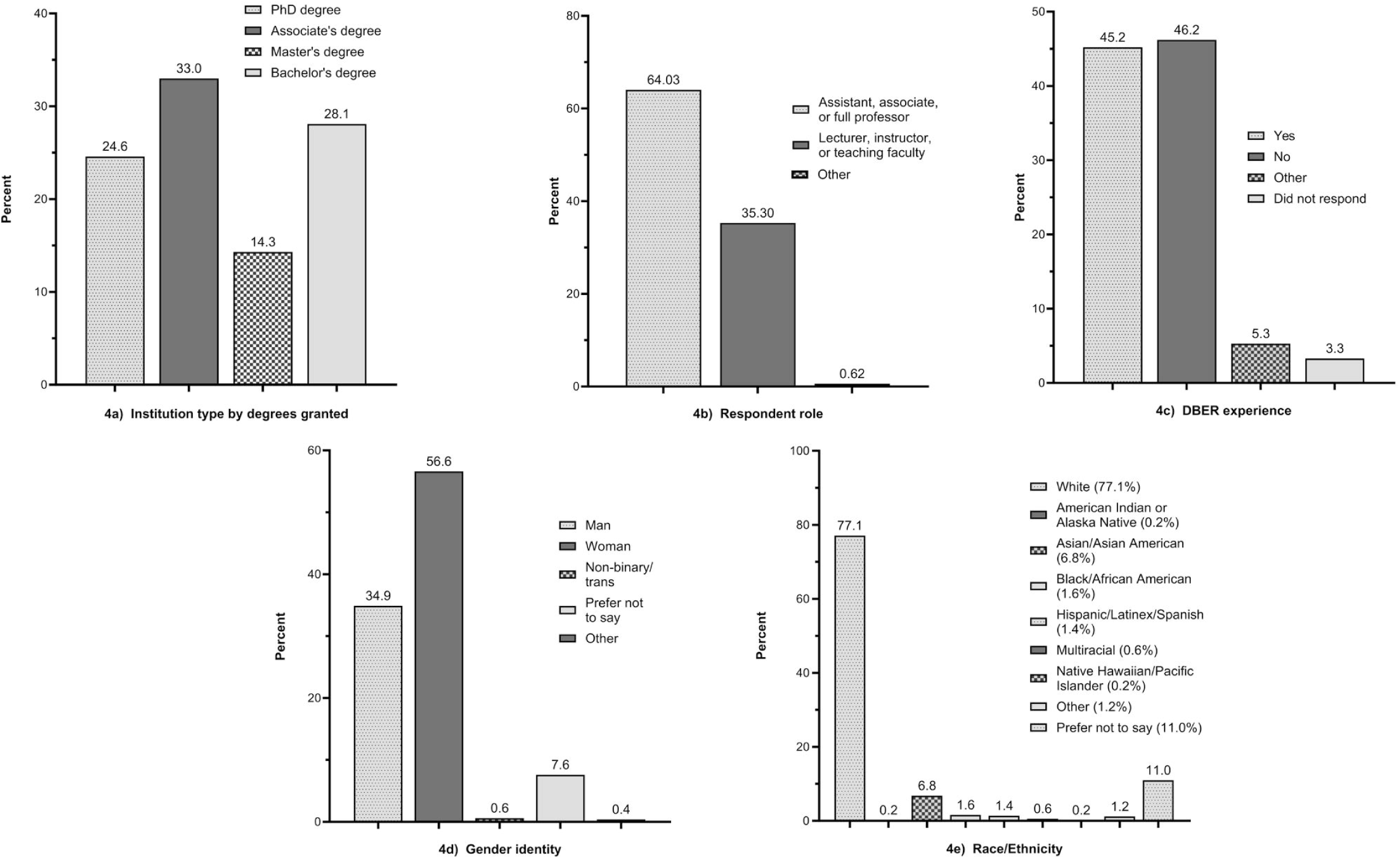
Self-reported demographics of LO evaluators.

### Binning the data into low, medium, or high support

S1 Data Set contains complete data on all 352 of the candidate LOs, including how many instructors evaluated it during the survey, what percentage of evaluators considered it essential, whether the LO requires LOCS or HOCS to master, and which *Vision and Change* Core Concepts and Competencies, BioCore Guide statements, and BioSkills Guide statements the LO aligns to.

Once this master dataset was assembled, we began data analysis by plotting the percentage of evaluators who had rated each LO as essential, creating a histogram representing responses for all 352 candidate LOs. Visual inspection of this graph revealed distinct break points in the data at 52.1% and 76.0% (S2 Fig). After confirming the existence of these breaks with Groups 2 and 3, we divided the data into three bins. We labeled LOs that had been ranked as essential by 52.1% or fewer respondents as “Low,” those considered essential by 52.2% - 76.0% of instructors as “Medium,” and those that were rated essential by 76.1% or more of respondents as “High” (Fig 5).

**Fig 5.**
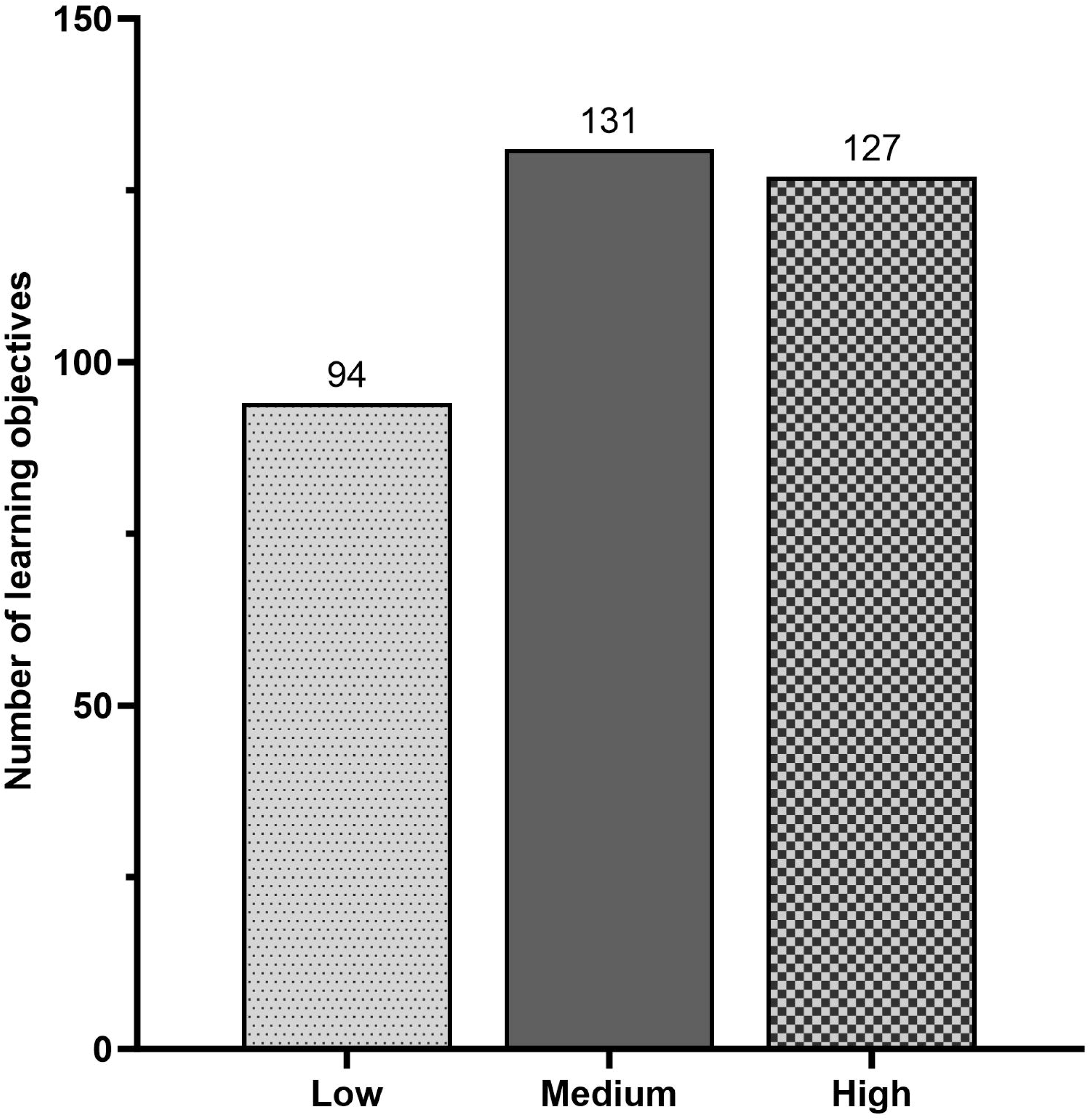
Summary of low, medium, and highly endorsed learning objectives.

Although survey participants were unaware of the LOCS or HOCS designation for the candidate LOs they evaluated, a striking pattern emerged when we separated LOCS from HOCs LOs and graphed the resulting data in the Low, Medium, and High bins. Evaluators were much more likely to rate an LO as essential if it represented a lower-order cognitive skill, and much more likely to rate an LO as non-essential if it represented a higher-order cognitive skill (Fig 6).

**Fig 6.**
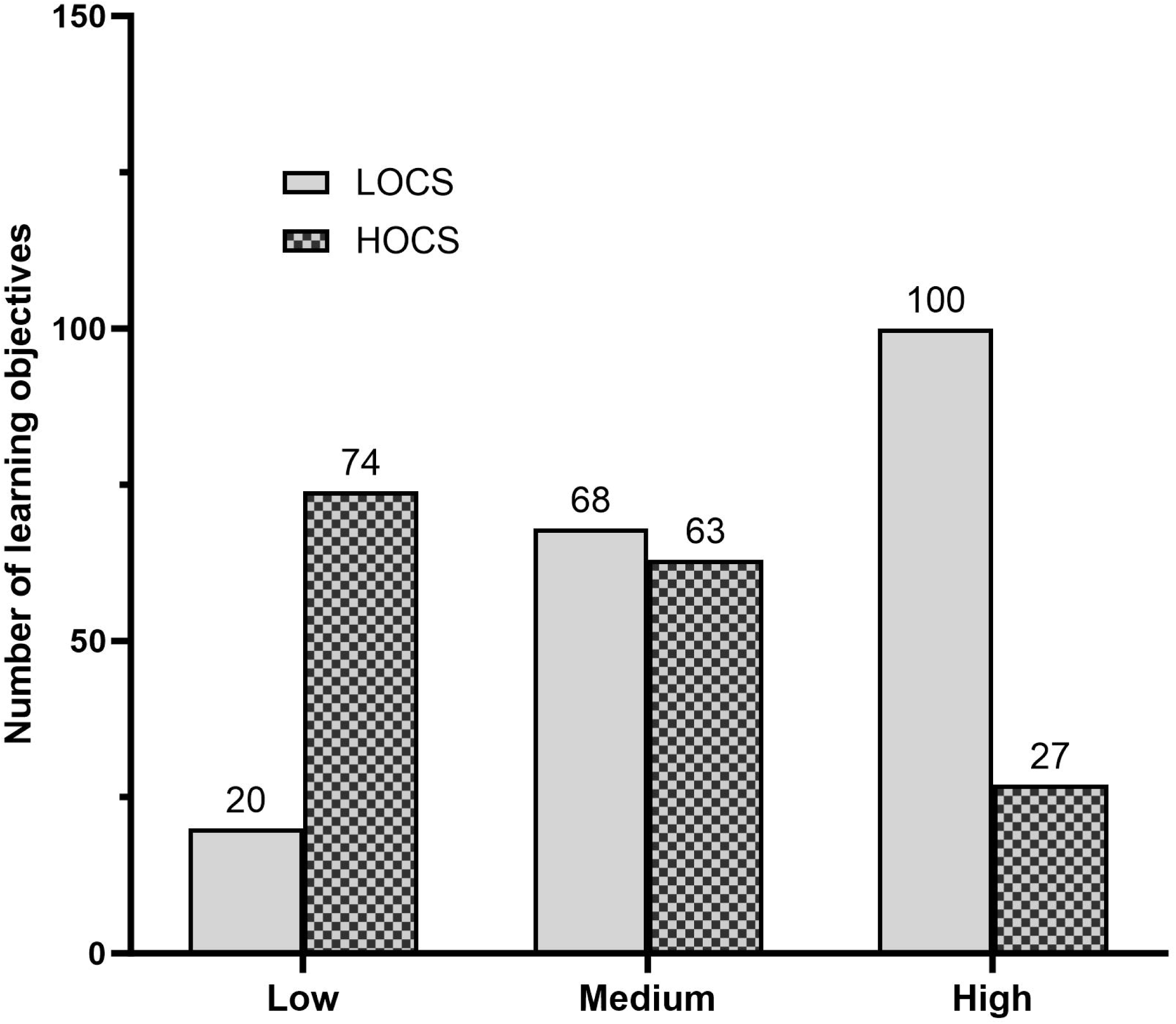
Summary of LOCS and HOCS separated by level of endorsement (low, medium, or high.)

This result was independent of topic area. When we graphed the data from each unit separately, the same pattern held across the majority of the units (S5 Fig).

### Recommending a set of LOs that are essential for majors biology

This project’s goal was to develop a set of lesson level LOs for a one-year introductory biology sequence that was consistent with programmatic goals articulated in the *Vision and Change* report and that have been endorsed as essential by a large, national sample of instructors. To support this goal, we recommend that faculty use all of the LOCS LOs that received high endorsement based on percent-essential ratings, and all of the HOCS LOs that received either a medium or high endorsement of percent-essential ratings. Following these guidelines, we recommend a total of 163 of the 352 candidate LOs as the core LOs for introductory biology for majors courses (Table 1).

**Table 1.** Recommended core LOs for introductory biology for majors courses.

| **Biochemistry of life recommended learning objectives** |  |  |
| --- | --- | --- |
| **Topic** | **Locs** | **Hocs** |
| Water | Draw the structure of several water molecules that are interacting and indicate 1) the electron distributions in each covalent bond, 2) the partial charges on each atom, and 3) each hydrogen bond. | None (appropriate HOCS LOs require biochemistry concepts beyond the scope of intro bio) |
| Water | Compare hydrogen bonds and covalent bonds in terms of the mechanisms and strength of attraction between the atoms involved. | Explain the relationship between hydrogen bonding and phenomena such as sweating, moderate coastal climates, and the oceans' response to global warming. |
| Water | Define the terms acid, base, and pH. Sketch the pH scale and note where on the scale you find strong acids, strong bases, and water | None (appropriate HOCS LOs require concepts in acid-base chemistry that are beyond the scope of intro bio) |
| Proteins | Label the four components of an amino acid and explain the role of each in terms of how the molecule functions in a protein. | Predict whether the R-group on an amino acid that you haven't seen before will 1) interact with water, and 2) act as an acid (proton donor) or base (proton acceptor). |
| Proteins | Describe each of the four levels of protein structure and explain how each influences the protein's final size, shape, and chemical properties | Label elements of primary, secondary, tertiary, and quaternary structure on a model of a protein that you haven't seen before |
| Proteins | Compare which bonds are responsible for producing a protein's 1) primary structure, 2) secondary structure (alpha-helices and beta-pleated sheets), and 3) tertiary structure | None (appropriate HOCS LOs are found in subsequent units that explore protein function in depth) |
| Proteins | Describe at least three functions that proteins serve in cells | None (appropriate HOCS LOs are found in subsequent units that explore protein function in depth) |
| Nucleic Acids | Define complementary base pairing, and explain its connection to the observation that DNA strands are antiparallel | Use the pairing rules to 1) explain the observation that in DNA, %A = %T and %G = %C, 2) predict the sequence of a complementary strand of DNA when given one strand, and 3) calculate the |
|  |  | percentage of each base in a DNA molecule when given the percentage of one base |
| Lipids | Use drawings, models, or other representations to compare the structures of fats, phospholipids, and steroids | Given a structural model of a lipid you've never seen before, 1) identify it as a fat, phospholipid, or steroid, 2) determine if it is saturated or unsaturated, and 3) predict its function in the cell |
| Lipids | Label the hydrophilic head and hydrophobic tails on a drawing of a phospholipid, then make drawings that include water molecules to explain how phospholipids spontaneously form bilayers in water | Given several models of membranes, predict how differences in phospholipid composition and cholesterol content will affect their relative fluidity and permeability, and explain your reasoning. |
| Membrane Structure and Transport | Draw a cell membrane and label integral and peripheral proteins, carbohydrate components, and lipid components. | None (appropriate HOCS LOs require concepts [e.g., diversity in membrane biochemistry] beyond the scope of intro bio) |
| Membrane Structure and Transport | Compare the processes of diffusion, osmosis, and facilitated diffusion, and provide biological examples that illustrate each process. | None, because appropriate HOCS LOs are found in subsequent units that explore movement of ions and molecules in depth. |
| Membrane Structure and Transport | Explain why ions and polar molecules do not move across plasma membranes efficiently without a transport protein. | Given several ions and molecules, predict the relative rates at which they will cross a plasma membrane in the absence of membrane proteins. Explain your reasoning. |
| Membrane Structure and Transport | Define passive and active transport and explain the role of channels, carriers, and pumps in transport. | None (appropriate HOCS LOs are found in subsequent units that explore membrane protein function in depth) |
| Comparing The Major Classes of Biological Molecules | Compare the monomer subunit, bond responsible for polymerization, and important biological function(s) observed in proteins, nucleic acids, and carbohydrates | Analyze how the structure of biological molecules impacts their function, including explaining the connections among the following three statements: 1) amino acids are much more diverse in structure and chemical properties than nucleotides, 2) in terms of diversity in shape and chemical properties, proteins > RNA > DNA, and 3) in terms of diversity in function, proteins > RNA > DNA. |
| Comparing The Major Classes of Biological Molecules | Compare the primary, secondary, and tertiary structures of proteins, RNA, and DNA |  |
| **Cells recommended learning objectives** |  |  |
| **Topic** | **Locs** | **Hocs** |
| Cell Structure and Function | Compare key elements of prokaryotic versus eukaryotic cell structure. | Propose hypotheses to explain 1) the adaptive significance of organelles (the advantages and disadvantages of having membrane-bound structures inside cells), and 2) why organelles are more common in eukaryotes than bacteria and archaea. |
| Cell Structure and Function | Compare key elements of plant versus animal cell structure. | None - HOCS received low endorsement during evaluation phase or beyond the scope of intro bio. |
| Cell Structure and Function | None (appropriate LOCS tasks--recalling the structure and function of cell parts--are embedded in the HOCS LO) | Predict what would happen to a cell if a particular organelle or structure was altered in a specified manner. |
| Cell Structure and Function |  | Predict 1) the function of a cell, given a drawing, micrograph, or description of its structure and organelle content, and 2) the structure of a cell, given information on its function. In each case, explain your reasoning. |
| Cell Structure and Function |  | Predict whether photosynthesis and/or cellular respiration will occur in a specific plant or animal cell, based on information about the cell's structure and function. |
| Cell Structure and Function | Make a flow chart showing how proteins are processed and packaged or unpackaged as they move from ribosomes to the interior of the rough ER to Golgi to motor proteins to their destination. | Predict what would happen to a particular protein or overall cell function if a specified element or process in the endomembrane system were altered. |
| Cell Structure and Function | Compare the structure and function of microtubules, actin filaments (microfilaments), and intermediate filaments. | None - HOCS received low endorsement during evaluation phase or beyond the scope of intro bio. |
| Cell Cycle/Mitosis | None (appropriate LOCS tasks--defining terms--are embedded in the HOCS LO) | Given a micrograph or drawing of a cell you've never seen before, label the chromosomes, chromatids, sister chromatids, and homologous chromosomes, if present, and determine the haploid number and ploidy. |
| Cell Cycle/Mitosis | Explain why chromosome replication has to occur before mitosis, in interphase. | Given a micrograph or drawing of a cell you've never seen before that is undergoing mitosis, explain what is currently happening to the chromosomes. |
| Cell Cycle/Mitosis | Given a labeled drawing showing the phases of mitosis, explain what is happening to the chromosomes and how it helps ensure that each daughter cell gets a complete and identical set. | None - HOCS received low endorsement during evaluation phase or beyond the scope of intro bio. |
| Cell Cycle/Mitosis | Diagram the sequence of stages in the eukaryotic cell cycle (M, G1, S, and G2) and label the major event or events that occur in each. | Predict the consequences of altering a given stage (M, G1, S, and G2) in the cell cycle in terms of the cell's structure or fate. |
| Respiration | Make a chart summarizing the inputs and outputs of glycolysis, pyruvate processing, the citric acid cycle, and oxidative phosphorylation, using NADH, FADH<sub>2</sub>, Glucose, Acetyl CoA, Pyruvate, O<sub>2</sub>, CO<sub>2</sub>, H<sup>+</sup> gradients, and ATP. Using the chart, explain how energy is transferred or transformed in each stage. | Predict the possible consequences if a step in the glucose oxidation (cellular respiration) pathway is altered. |
| Respiration | Explain how cells use fermentation pathways to obtain energy from glucose in the absence of oxygen. | None (appropriate HOCS LOs require biochemistry concepts [e.g., experimental evidence] beyond the scope of intro bio. |
| Respiration | None (appropriate LOCS tasks--identifying parts of the electron transport chain or ATP synthase--are embedded in the HOCS LO) | Predict the effects of altering specific parts of the electron transport chain or ATP synthase. |
| Photosynthesis | Given the overall formula for photosynthesis, label the atoms that lose electrons, the atoms that gain electrons, the molecules that are reduced, and the molecules that are oxidized. Explain the impacts of | Explain to an 8-year-old how the CO<sub>2</sub> in "weightless" air is the source of mass in a redwood tree. |
|  | these changes on the energy available in each molecule. |  |
| Photosynthesis | Make a chart summarizing the inputs and outputs of PSI, PSII, and the Calvin cycle using NADPH, Glucose, H<sub>2</sub>O, O<sub>2</sub>, CO<sub>2</sub>, H<sup>+</sup> gradients, and ATP. Using this chart, explain the energy transformations that occur and the role of rubisco. | Predict the possible consequences for the production of ATP and NADPH if a component or process in the photosynthesis pathway is altered. |
| Enzymes/Energetics | None (appropriate LOCS tasks [e.g., defining free energy, exergonic, endergonic] are embedded in the HOCS LO) | Given a graph showing how free energy changes over the course of a chemical reaction, 1) label the sections representing the reactants, activation energy, and products, 2) explain why energy peaks during the transition state, and 3) determine whether the reaction is exergonic or endergonic. |
| Enzymes/Energetics | Explain 1) why "active site" is an appropriate term, 2) the mechanisms responsible for the observation that enzymes lower activation energies, 3) why most enzymes catalyze one specific reaction, and 4) why enzymes increase reaction rates but do not make endergonic reactions exergonic. | None (appropriate HOCS LOs require biochemistry concepts [e.g., experimental evidence] beyond the scope of intro bio. |
| Enzymes/Energetics | None, (appropriate LOCS tasks are generalized graph-reading skills) | Interpret graphs of reaction rate versus pH, temperature, and degree of substrate saturation for a given enzyme. Based on your analysis, predict the nature of the cell's normal environment in nature. |
| Enzymes/Energetics | Explain 1) the general role of ATP in the cell, 2) what it means to say that two chemical reactions are coupled, and 3) why a large change in free energy level occurs when an enzyme or substrate is phosphorylated. (Recall that phosphorylation adds 3 tightly packed negative charges.) | Given graphs showing how free energy changes over the course of a chemical reaction, predict whether two specific reactions can be successfully coupled. |
| Overview | Given the summary reactions for photosynthesis and respiration, | None (appropriate HOCS LOs require biochemistry concepts [e.g., |

|  | compare 1) the reactants and products of each process, and 2) the energy transformations that occur. | experimental evidence] beyond the scope of intro bio. |
| --- | --- | --- |
| **Genetics recommended learning objectives** |  |  |
| **Topic** | **Locs** | **Hocs** |
| Meiosis | Explain the differences between somatic cells and germ cells. Describe the outcomes of cell division between these two categories of cells. | None (appropriate HOCS tasks [e.g., analyzing relationships in plants and other groups] are beyond the scope of intro bio) |
| Meiosis | Differentiate between the genetic information held on two homologous chromosomes, two nonhomologous chromosomes, two sister chromatids, and two non-sister chromatids. | Given a micrograph or drawing of a cell you've never seen before, label chromosomes, chromatids, sister chromatids, and homologous chromosomes, if present, and determine the haploid number and ploidy of the cell. |
| Meiosis | None (appropriate LOCS tasks [e.g., listing the issues involved] are embedded in the HOCS LO) | Given a micrograph or drawing of a cell you've never seen before that is undergoing meiosis, explain what is currently happening to the chromosomes. |
| Meiosis | Explain why the segregation of homologous chromosomes in meiosis I leads to a reduction in ploidy. | Given a specific error in meiosis, predict the haploid genotypes that result and discuss the consequences for offspring. |
| Meiosis | Explain why no two haploid cells that result from meiosis are alike in terms of genotype and why this is important in terms of offspring fitness. | None (appropriate HOCS tasks are beyond the scope of intro bio) |
| Mutations/Cancer | Rank the following mutations in terms of greatest to least impact on the structure and function of genes and gene products: missense (change amino acids), nonsense (change to "stop"), frameshift (change reading frame), and silent (no change in the product). Explain your reasoning. | None (appropriate HOCS tasks [e.g., analyzing examples in real data] are beyond the scope of intro bio. |
| Mutations/Cancer | Defend the statement "mutation is the ultimate source of genetic variation," and explain why | None (appropriate HOCS tasks [e.g., analyzing variation in mutation rate among species, or |
|  | mutation is random with respect to its impact on an individual's fitness. | codon bias] are beyond the scope of intro bio. |
| Mutations/Cancer | Explain why cancer is 1) associated with mutations that regulate the cell cycle, and 2) more common in older than younger people. | None (appropriate HOCS tasks [e.g., analyzing mutations associated with a specific cancer] are beyond the scope of intro bio) |
| Patterns of Inheritance | Label which elements in a Punnett square represent the genotypes of egg, sperm, and offspring. Explain how you can determine the frequency of each egg and sperm genotype and how you can use this information to calculate the frequencies of offspring genotypes and phenotypes. | Given any pair parental genotypes and information on the alleles present, use a Punnett square to complete a genetic cross. Identify the genotypes and phenotypes of offspring and calculate their predicted frequencies. Note that the genes involved may be autosomal, X-linked, linked, or unlinked and that the alleles involved may be dominant, recessive, or co-dominant. |
| Patterns of Inheritance |  | Given information on parental and offspring phenotypes, determine whether the alleles involved are 1) dominant, recessive, or codominant, 2) autosomal or X-linked, and 3) linked or unlinked. |
| Patterns of Inheritance | Define polygenic inheritance and explain why it produces traits with a continuous variation. | None (appropriate HOCS tasks are beyond the scope of intro bio) |
| Patterns of Inheritance | Using a drawing that shows the phases of meiosis, label the events that explain Mendel's principles of segregation and independent assortment. Add drawings to show how independent assortment can generate genetic variation in offspring. In each case, explain your reasoning. | None (appropriate HOCS tasks [e.g., relating linkage and recombination as exceptions] are beyond the scope of intro bio) |
| Patterns of Inheritance | On a pedigree, label 1) males and females, 2) affected and unaffected individuals, and 3) generations. | Based on the data in a pedigree, predict 1) whether the trait in question is autosomal or sex-linked and 2) which alleles are dominant and recessive. |
| DNA Replication | Describe the function of major components of the replisome: helicase, topoisomerase, DNA polymerase, DNA ligase, and primase. | None (appropriate HOCS tasks [e.g., analyzing the structure of each component or molecular mechanisms involved] are beyond the scope of intro bio) |
| DNA Replication | Use a drawing that you create to explain the statement: "A newly synthesized DNA strand is half old and half new." | None (appropriate HOCS tasks are of historical interest only [e.g., Meselson-Stahl experiment] or beyond the scope of intro bio [e.g., self-replicating RNAs]) |
| DNA Replication | Given a diagram of a DNA molecule during replication, label the following: the origin of replication, directions of replication, replication fork, the leading strand, and lagging strands and their polarities, and the replisome. | Using everyday objects like twine or fabric strips, create a model of DNA replication and use it to explain 1) why lagging strand synthesis is an appropriate name, and 2) why Okazaki fragments occur. |
| DNA Replication | Explain how DNA damage and/or mismatches are detected and repaired. | None (appropriate HOCS tasks are beyond the scope of intro bio) |
| Information Processing | Make a flow chart summarizing the flow of information in cells from gene to protein. Label arrows connecting mRNA, DNA, and proteins, and explain what each arrow represents. | Add elements to your central dogma model that represent "exceptions" such as 1) production of rRNA, tRNA, and "other RNAs", 2) DNA replication, and 3) the action of an enzyme called reverse transcriptase, which catalyzes the synthesis of DNA from an RNA template. |
| Information Processing | Explain how the genetic code relates transcription to translation and why it is considered redundant. | None (appropriate HOCS tasks [e.g., analyzing exceptions to the canonical genetic code] are beyond the scope of intro bio) |
| Information Processing | On diagrams of transcription initiation and transcription elongation, label the template and coding strands, initiation complex, promoter site, RNA polymerase, ribonucleotides, the direction of RNA polymerase movement, and direction of RNA synthesis. | Given a specific change in a DNA coding strand or a specific error in transcription or translation, predict the consequences for the gene product. |
| Information Processing | On diagrams of translation initiation, translation elongation, and translation termination, label the small and large ribosomal subunits, mRNA, tRNA, rRNA, reading frame, start codon, stop codon, release factor, and tRNA binding sites (E, A, and P). Circle and label the locations where | Use a copy of the genetic code to predict the sequence of the amino acids produced from a given mRNA or double-stranded DNA fragment. Identify the start and stop codon. |
|  | codon- anticodon recognition and peptide bond formation occur. |  |
| Information Processing | Compare the structure, chemical composition, location, and function of DNA with RNA. | None (appropriate HOCS tasks [e.g., analyzing the RNA world hypothesis for origin of life] are beyond the scope of intro bio) |
| Control of Gene Expression | Explain how negative and positive control over transcription regulates the activity of a given gene or operon. | None (appropriate HOCS tasks are beyond the scope of intro bio) |
| Biotechnology | None (appropriate LOCS tasks are embedded in the HOCS LO) | Given an image of a gel, predict the relative size of protein or DNA samples and interpret the presence or absence of a known protein or region of DNA (such as a PCR product). |
| **Evolution recommended learning objectives** |  |  |
| **Topics** | **Locs** | **Hocs** |
| Natural Selection | Define adaptation, fitness, evolution, and theory. For each term, explain how its use in science differs from its use in everyday English. | None (appropriate HOCS tasks [e.g., analyzing controversies over teaching evolution or evaluating alternative theories for the origin of species] are beyond the scope of intro bio) |
| Natural Selection | Explain why the fossil record and genetic and structural homologies provide evidence for evolution. | None (appropriate HOCS tasks [e.g., relating genetic to structural homologies] are beyond the scope of intro bio) |
| Natural Selection | Explain the connection between mutation and heritable variation in traits, and how selection on this variation can lead to changes in allele frequencies. | Analyze data or design an experiment in the lab, greenhouse, or field on evolution by natural selection. Identify the model organism, treatment, control conditions, and outcome variable measured. Interpret the outcomes, or graph the predicted outcomes, for both treatments over many generations. |
|  |  | Using specific examples, explain why evolution by natural selection is neither random nor progressive, and why adaptations are not 'perfect'. |
| Other Evolutionary Processes | Define genetic drift and describe how it influences allele frequencies. Explain why it is more important in small populations than in large populations and why it eventually leads to fixation or loss of alleles. Provide examples of events or processes that cause drift. | None (appropriate HOCS tasks are beyond the scope of intro bio) |
| Other Evolutionary Processes | Define gene flow and describe how it impacts allele frequencies in the source and recipient population. | None (appropriate HOCS tasks are beyond the scope of intro bio) |
| Other Evolutionary Processes | Defend the statement "mutation is the ultimate source of genetic variation," and explain why mutation is random with respect to its impact on an individual's fitness. | None (appropriate HOCS tasks [e.g., analyzing variation in mutation rate among species or codon bias] are beyond the scope of intro bio) |
| Other Evolutionary Processes | Explain 1) why the Hardy-Weinberg principle provides a null model for evolution, and 2) why natural selection, genetic drift, gene flow, mutation, and/or non-random mating can each produce genotype frequencies different from those expected under the Hardy-Weinberg principle. | For a gene with a given number of alleles, identify the genotype frequencies expected under the Hardy-Weinberg principle. |
| Tree Thinking | None, (appropriate LOCS tasks [e.g., defining terms such as root and node] are embedded in the HOCS LO) | Given a phylogenetic tree, 1) label the root, nodes, branches, and tips; 2) identify the type of taxon at the tips (e.g., species or larger group), 3) circle and label several monophyletic groups, at least two of which are nested, 4) state at least one synapomorphy that identifies each of the circled monophyletic groups, and 5) find the most recent common ancestor of any two given taxa. |
| Tree Thinking | None (appropriate LOCS tasks [e.g., explaining the use of parsimony as a decision criterion] are embedded in the HOCS LO) | Given a phylogeny and information about a trait in species included, mark the tree to indicate where changes in the trait occurred. |
| Speciation | Explain 1) why genetic isolation and genetic divergence lead to speciation, and 2) the roles that gene flow, natural selection, genetic drift, and mutation can play in | Analyze data or design an experiment on how two populations may be evolving into distinct species |
|  | speciation. |  |
| History of Life | Define adaptive radiation and use examples to explain how ecological opportunities and/or key innovations can cause them. | None (appropriate HOCS tasks are beyond the scope of intro bio) |
| Synthesis | Explain how gene flow, inbreeding, and genetic drift may positively or negatively affect endangered species that live in isolated (fragmented) habitats. | None (appropriate HOCS tasks are beyond the scope of intro bio) |
| **Biodiversity of life recommended learning objectives** |  |  |
| **Topic** | **Locs** | **Hocs** |
| Tree of Life | None (appropriate LOCS tasks are embedded in the HOCS LO) | Classify an organism that you've never heard of before, by domain, when given a set of its characteristics. Explain your reasoning. |
| Tree of Life | None (appropriate LOCS tasks [e.g., recalling key traits associated with major taxa] are embedded in the HOCS LO) | Given a set of traits, identify a species that you've never encountered before as a virus, "protist," plant, fungus, or animal, and explain your reasoning |
| Bacteria and Archaea | Describe at least three of the beneficial and harmful roles that bacteria and archaea play in the lives of humans and the biosphere as a whole. | Explain why antibiotics specifically kill bacteria instead of the host, and why antibiotics that kill many different species of bacteria tend to cause side effects for the host. |
| Eukaryotic Radiation Protists | Describe the key characteristics of the common ancestor of all eukaryotes living today. | Using information provided on the traits of several major lineages, construct or label a tree of the eukaryotic radiation to show where key synapomorphies arose. |
| Eukaryotic Radiation | Explain the process of endosymbiosis and using your knowledge or information provided on a specific example, describe the costs and benefits to each organism involved. | None (appropriate HOCS tasks are beyond the scope of intro bio) |
| Plants | None (appropriate LOCS tasks [e.g., recalling facts on morphology] are embedded in the HOCS LO) | Explain why the following sources of evidence support the hypothesis that land plants evolved from freshwater green algae: 1) |
|  |  | morphological data, 2) the fossil record, and 3) the phylogenetic tree. |
| Plants | On a tree showing the relationships among major lineages of green algae and land plants, circle the most species-rich tip and label where the following synapomorphies arose: vascular tissue, flowers, stomata, cuticle, seeds, cellulose cell walls, pores, chloroplasts. Explain the adaptive significance of each synapomorphy. | None (appropriate HOCS tasks [e.g., explaining trait loss or origin of key traits] are beyond the scope of intro bio) |
| Plants | Summarize the major features of alternation of generations. | None (appropriate HOCS tasks are beyond the scope of intro bio) |
| Plants | Describe how plants influence the environment in terms of 1) atmospheric composition, 2) food availability in terrestrial environments, 3) soil formation, and 4) the quality and abundance of water in soils, marshes, and streams. | Predict the impact of extensive deforestation or other changes in plant communities in terms of 1) atmospheric composition, 2) food availability in terrestrial environments, 3) soil formation, and/or 4) the quality and abundance of water in soils, marshes, and streams. |
| Fungi | Use a drawing or other graphic to explain the mutually beneficial relationships between fungi and plants. | None (appropriate HOCS tasks are beyond the scope of intro bio) |
| Viruses | Explain why viruses are considered obligate intracellular parasites, and why viral diseases are difficult to treat with drugs. | State the arguments for why viruses could be considered alive or not alive, then state and defend your opinion on this question. |
| Viruses | Using a diagram, explain how the following events in a virus' life cycle occur: enter a host cell, produce viral proteins, replicate viral genome, assemble new virions, exit host cell, transmission to a new host. | None (appropriate HOCS tasks are beyond the scope of intro bio) |
| Animals | On a tree that shows the major lineages of animals, label where the following synapomorphies arose: bilateral symmetry, cephalization, multicellularity, movement via contractile proteins. Explain each synapomorphy's adaptive significance. Identify the two most | Using information that you gather or are provided, characterize the following in a particular major animal lineage: 1) body plan, 2) reproductive systems, 3) sensory organs, 4) feeding strategies, and 5) ecological role. |

|  | species-rich lineages. |  |
| --- | --- | --- |
| **Plant and animal physiology recommended learning objectives** |  |  |
| **Topic** | **Locs** | **Hocs** |
| Plant and Animal Physiology | None (appropriate LOCS tasks [e.g., listing the components of the system in question] are embedded in the HOCS LO) | Analyze the relationships among the cells, tissues, and/or organs involved in a given physiological system, including how their structures correlate with their functions and how they interact in terms of function. |
| Flow Down Gradients | Explain how and why ions and molecules move in response to concentration gradients. | None (appropriate HOCS tasks are beyond the scope of intro bio) |
| Flow Down Gradients | Explain how ions move in response to electrical potential gradients, and why. | Based on the information you are given about a specific cell, predict how it will be affected by ion movements that occur in response to a change in electrical potential gradients. |
| Cell to Cell Communication | Compare the structure and solubility of peptide, steroid, and amine messengers, then compare how they are transported in the blood and enter cells. | None (appropriate HOCS tasks [e.g., analyzing hormone-hormone interactions] are beyond the scope of intro bio) |
| Cell to Cell Communication | Explain the relationship between a chemical messenger and its receptor, and why only certain cells respond to a specific messenger. | Predict how a given change in a receptor will alter a cell's response. |
| Cell to Cell Communication | Explain why chemical messengers can elicit a large response even though they are present at very low concentrations. | None (appropriate HOCS tasks are beyond the scope of intro bio) |
| Electrical Signaling | Draw a generalized version of a neuron. Label the dendrites, cell body, an axon and describe the function of each. | None (appropriate HOCS tasks [e.g., analyzing change in cell morphology over time or consequences of damage or disease] are beyond the scope of intro bio.) |
| Electrical Signaling | Explain 1) what it means to say that an action potential is an all-or-none event, and 2) how specific information can be represented by the frequency and source of action | Given data on the action potentials received by a cell predict whether the cell will fire an action potential in response. |
|  | potentials received by a cell. |  |
| Electrical Signaling | Explain how changes in membrane potential that occur during an action potential depend on the activity of voltage-gated channels, and how an action potential propagates along a cell. | None (appropriate HOCS tasks are beyond the scope of intro bio) |
| Electrical Signaling | Explain how the information in an action potential is transduced at a synapse and then transmitted to a post-synaptic cell. | Given information on a specific neurotoxin, predict how it will impact the transmission of action potentials. |
| Homeostasis | None (appropriate LOCS tasks [e.g., listing the components of the system in question] are embedded in the HOCS LO) | For a given homeostatic system, predict how a given change in one or more of its components will change the activity of the system. |
| Homeostasis | For a given homeostatically regulated property, explain how positive and negative feedback maintains it at a normal range and why that is important to the organism's fitness. | None (appropriate HOCS tasks are beyond the scope of intro bio) |
| Plant Specific Physiology | In a drawing or photograph of a plant you have never seen before, identify roots, stems, and leaves and explain their overall function. | Given drawings or photographs and information on the habitat occupied by a plant you've never seen before, suggest hypotheses for the adaptive (functional) significance of its root, stem, and leaf structures. |
| Sensing and Responding | Describe how stomata (paired guard cells) allow for CO<sub>2</sub> and O<sub>2</sub> exchange and explain how and why they change shape in response to changes in the environment. | Under given light, water, or temperature conditions, predict adaptations or developmental responses in stomata and other cells and structures that optimize the rate of photosynthesis and minimize the rate of water loss. |
| Growth and Reproduction | Given a diagram of a familiar plant lifecycle, label when and where mitosis, meiosis, and fertilization occur, and identify haploid and diploid phases. | Given a diagram of the life cycle of a plant you have never seen before, label when and where meiosis and fertilization occur and identify haploid and diploid phases. |
| Matter and Energy Flow | Create a diagram, drawing, or model to communicate how carbon is assimilated into organic compounds in plants and relate this to the flow of energy and the synthesis of molecules required for | Explain how the CO<sub>2</sub> in "weightless" air is the source of mass in a redwood tree to an 8-year-old. |
|  | maintenance and growth. |  |
| Matter and Energy Flow | Create a drawing or other model to explain how inorganic nutrients are obtained from soil, either directly or via associations with mycorrhizal fungi. | None (appropriate HOCS tasks [e.g., analyzing changes during development or the consequences of disease, damage, or domestication] are beyond the scope of intro bio) |
| Matter and Energy Flow | Compare the structure and function of the cells and tissues involved in sugar transport (phloem) versus water transport (xylem). | Compare the mechanisms responsible for the long-distance transport of sugars versus water and nutrients. |
| **Ecology recommended learning objectives** |  |  |
| **Topic** | **Locs** | **Hocs** |
| Introduction | Explain the difference between abiotic and biotic factors in an ecosystem and provide examples of how these factors can affect the communities present. | None (appropriate HOCS tasks are beyond the scope of intro bio) |
| Introduction | Create a diagram explaining the nested relationships among populations, species, communities, and ecosystems. | None (appropriate HOCS tasks [e.g., explaining the limitations or dynamic aspects of these categories] are beyond the scope of intro bio) |
| Populations | Given information presented as graphs, tables, or equations, describe how the size of a population has changed over time. | None (appropriate HOCS tasks are beyond the scope of intro bio) |
| Species Interactions and Community Ecology | Explain why life-history traits, competition, predation, parasitism, mutualism, and niche breadth (abiotic factors) impact the distribution and abundance of a given species. | None (appropriate HOCS tasks [e.g., spatial-temporal dynamics, conservation implications, multiple and simultaneous interactions] are beyond the scope of intro bio) |
| Species Interactions and Community Ecology | Explain why life-history traits, competition, predation, parasitism, mutualism, and niche breadth (abiotic factors) impact which species are found in the same community. | Analyze the consequences of human activities changing a community's distribution from the continuous occupation of a broad area to occupying small, disconnected patches within the same area. |
| Species Interactions and Community | None, (appropriate LOCS tasks are embedded in the HOCS LO) | Given a food web or other information on relationships among |
| Ecology |  | species in that community, predict the consequences of removing an apex ("top") predator, introducing a specific disease, or other specified changes occurring. |
| Matter and Energy in Ecosystems | Describe the major events in the global cycling of water, carbon, and nitrogen. | None (appropriate HOCS tasks [e.g., analyzing implications of changing specific elements of a cycle] are beyond the scope of intro bio) |
| Matter and Energy in Ecosystems | Explain how human activities have affected global nutrient cycles. | None (appropriate HOCS tasks are beyond the scope of intro bio) |
| Matter and Energy in Ecosystems | Explain why there is typically less biomass and fewer individuals and species at the top of a food chain than at the bottom. | Predict the consequences of changes in primary production due to perturbations such as drought, fire, flood, extreme temperatures, or nutrient influx or loss. |
| Climate Change | Interpret graphs that show changes in atmospheric CO<sub>2</sub> over the past 200 years and the past 10 years. | None (appropriate HOCS tasks are beyond the scope of intro bio) |
| Climate Change | Explain how human activities impact climate change by disrupting the global carbon cycle. | None (appropriate HOCS tasks [e.g., analyzing changes in conditions] are beyond the scope of intro bio) |
| Climate Change | Explain how climate warming leads to changes in the timing of seasonal events (phenology) and how changes in the phenology of one species can affect the ecology of other species. | None (appropriate HOCS tasks are beyond the scope of intro bio) |
| Biodiversity | Explain how biologists measure biodiversity, how species richness differs from species diversity, and why plant diversity is often used as an indicator of overall biodiversity. | Analyze data or design a study on current biodiversity at a site and changes in biodiversity over time. |
| Biodiversity | Explain how habitat loss, habitat fragmentation, climate change, over-harvesting, and invasive species affect biodiversity. | None (appropriate HOCS tasks are beyond the scope of intro bio) |

A total of 163 core learning objectives (LOs) are recommended for introductory biology courses for majors, comprised of both LOCS and HOCS LOs that received high endorsement based percent-essential ratings.

To evaluate how this total of 163 would be distributed over a yearlong course, we assumed that instructors would design their course using an average of three lesson level LOs per class session and followed Heil, et al.’s (2023) research indicating an average of 72 class sessions (“lectures”), exclusive of exams and holidays, in a two-semester course. This total of 216 LOs (three per session x 72 class sessions) could then be partitioned into a core set of 163 (∼75%) represented by the national endorsement reported in this paper and an additional 53 (∼25%) developed by individual instructors or teaching teams to reflect LOs germane to their program and student population.

Over 76% of the recommended learning objectives listed in Table 1 are included in LOCS- HOCS pairings. Unpaired LOs occurred when the natural LOCS LO for a HOCS LO would have consisted of a simple list of terms or other memorized information so was considered implicit; or the HOCS that was proposed was either deemed beyond the scope of introductory biology during the development phase or received low endorsement during the evaluation phase.

## Discussion

There are at least three major and interrelated justifications for using lesson level LOs that are well-written and appropriately scaffolded and framed during instruction [5]:

1. LOs are essential to backward design–widely considered to be the gold standard in course design.
2. When directed or guided by instructors, LOs give students a reliable and efficient way to focus their study efforts–enough so that instructors should no longer hear questions stems like, “Do we have to know” or “Will X be on the test? And,
3. LOs directly address two salient characteristics of novice learners that have been characterized by cognitive scientists: an inability to distinguish more-important from less-important information, and an inability to see connections between topics or integrate information [4].

The set of 163 core LOs presented here represents national endorsement for essential content, concepts, and skills. They should support instructional teams that want to introduce or expand the use of LOs in courses while aligning instruction with the national consensus articulated by *Vision and Change* [9]. It should also support ongoing efforts to correct the misalignment between course-level learning goals and assessment items that has plagued life sciences education for years [22].

These LOs complete a longstanding effort to create a cohesive framework for undergraduate biology education, beginning with the seminal *Vision and Change* report, continuing with development of the BioCore Guide and BioSkills Guide, and now concluding with a nationally endorsed set of lesson-level LOs. Biology educators are now in the enviable position of having clear expectations articulated for what students should know and be able to do when students emerge from an introductory course sequence for majors (Fig 1).

A course designed around these 163 recommended LOs, even when supplemented by individual instructors or teaching teams with an additional 40-50 customized LOs, promises to be radically different from a course designed around textbook content. The leading textbooks for introductory majors biology are 1200-1500 pages long and contain more than 2110 boldfaced glossary words and phrases. Given an average number of class sessions in a canonical 3-quarter or 2-semester course sequence, instructors would need to insist that their students memorize 18- 29 new terms every single class session if the goal of the course is to “cover” the textbook. The 163 “core” LOs recommended by the community strip away many of these terms and many of the other details commonly found in textbooks. As a result, our hope is that this national endorsement for LOs will help liberate instructors from what we call “the tyranny of content,” allowing them to re-focus their course on only essential vocabulary and concepts. Most importantly, this allows instructors to devote much more time and effort to the analytical and professional skills that many programs state as their major learning goal [21,22] and that are required for successful careers related to the life sciences.

### The role of LOCS and HOCS LOs

Instructors were much more likely to rate an LO as essential if it addressed lower-order cognitive skills (LOCS) as opposed to higher-order cognitive skills (HOCS). This finding is consistent with literature showing that introductory courses for life sciences majors currently emphasize content coverage and memorization [21–23] However, this finding conflicts with repeated calls from education policy leaders for instructors to de-emphasize lower-order cognitive skills and increase emphasis on the analytical and other higher-order skills required for success as a life science professional [4,9].

The pattern of higher endorsement levels for LOCs and lower endorsement levels for HOCS reported in Fig 6 may, however, also reflect the underlying nature of LOCS and HOCS– specifically, the claim that Bloom’s taxonomy of learning is hierarchical [32]. In many or most cases, the ability to apply concepts in novel situations, analyze processes and data, synthesize information to create something new, and evaluate the quality of hypotheses or evidence–that is, to work at higher cognitive levels–depends on a fundamental understanding of the underlying facts and concepts.

In weighing the balance between using LOCS and HOCS LOs in introductory courses for majors, we follow the national consensus represented in *Vision and Change* and recommend a “both-and” strategy that recognizes the fundamental importance of LOCS but places a much stronger emphasis on HOCS than is traditional. To implement this approach, we used different criteria for the list of 163 recommended LOs by including HOCS that had a medium level of support as essential while requiring a higher level of support for LOCS. We claim that the relaxed criterion for including HOCS still represents an important national endorsement for what is essential to teach in introductory biology for majors, as medium support indicates affirmation by over 50% of respondents.

### Limitations of this study

The LOs that the community of introductory biology instructors developed and recommended through this work focuses on content: vocabulary, concepts, and skills. At the start of the study, we had also intended to develop and evaluate LOs that focused on affect. In particular, we were interested in aspects of emotional and psychological experience that are known to impact retention in STEM and that are considered fundamental to professional development. None of the over 3000 draft LOs that launched the project addressed these affect issues, however, and when we began the process of creating LOs on affect, during the project’s development phase, we realized that it was not feasible to write them at the lesson level. Stated another way, we did not see how affect LOs could be made granular enough to be compatible with the content- and skills-focused, lesson-level LOs developed in this study. Thus, instructors should be aware that LOs recommended here do not address course goals outside of the *Vision and Change* framework, including course goals relating to affect [9].

Another limitation of the study concerns the sample of biology instructors who evaluated the 352 candidate LOs. Although we broadcast the appeal for raters as widely as possible, respondents volunteered their time and thus represent a convenience sample and not a random sample of the entire community. Currently we lack the data on the demographics of introductory biology instructors in the U.S. required to assess how representative our sample was. It may be helpful to note, however, that the total of 77% white faculty who evaluated the candidate Los can be compared to the percentage of white STEM instructors at various institution types in the U.S.: 75% at non-minority-serving institutions, 70.5% at tribal institutions, 68% at Asian-American Native-American and Pacific Islander-serving institutions, 63% at Hispanic-serving institutions, and 27% at Historically Black Colleges and Universities [33].

### Future work

LOs are just one of three elements that must be in place to achieve an integrated course design. Although work on developing evidence-based teaching materials–including pre-class preparatory materials, active learning activities for use in class, and post-class assignments–has progressed rapidly [34], the third component of course-design, representing formative and summative assessment, is weak. Researchers and practitioners will have a great deal of work to do before instructors have reliable and valid assessment items that align with both the LOs published here and available teaching materials. This is particularly true of assessment items that are machine- gradable and yet require students to apply concepts, analyze processes or data, design experiments, or evaluate claims. It is a challenge to create authentic assessments that test HOCS LOs while providing timely feedback to students, but a challenge that the community needs to meet.

Another important frontier in research concerns student use of lesson-level LOs, and in particular how structured exercises and informal instructor talk can impact students [5]. Do students use LOs more effectively if instructors make the connection between specific LOs, teaching practices, and assessment items more transparent? Are there ways to structure student use of LOs during self-study and exam preparation? Does the use of LOs make students more efficient and thus effective in terms of using their study time? These questions are all unanswered.

Currently, life science educators also lack a national endorsement for course-level LOs that address aspects of affect and professional and career development relevant to prospective biology majors. If a follow-up study put these course-level statements in place, researchers and practitioners could begin implementing classroom practices and developing assessments capable of supporting student progress on these critically important elements of professional maturation and success.

Finally, this study should support future work on what we see as two extraordinarily challenging, but potentially extraordinarily rewarding, research endeavors:

1. Designing and executing a rigorous test of the backward design hypothesis–one that evaluates the claim that integrated course design leads to consistently better student outcomes; and
2. Developing learning progressions for particularly important LOs in the set recommended here, and then using an improved understanding of stages in the transition from novice to expert-level understanding to design improved teaching materials and assessment items.

## Conclusions

This is the first effort to develop a community-endorsed set of LOs that are comprehensive enough to serve as a core element in designing introductory biology courses for majors. It completes the “organizational chart” of community-endorsed learning goals that began with the publication of the *Vision and Change* report and continued with the development of the BioCore Guide and BioSkills Guide and should provide additional momentum to ongoing efforts to transform introductory biology courses for majors. Life science educators now have the most comprehensive, nationally endorsed course-design framework of any STEM discipline.

Integrating the LOs reported here should help focus courses in productive ways, free instructors from the pressure to “cover it all,” and create consistency and predictability within and across programs and institutions.

As the life sciences advance and as discipline-based education research yields new insights, however, the LOs published here should evolve in response. We look forward to revisions that benefit from new insights in the life sciences and the science of learning, and to efforts that use LOs to support student success.

## Acknowledgements

We thank the following people who were involved in the Development Phase of the grant (listed alphabetically):

Joel Abraham

Tessa Andrews

Norris Armstrong

Holly Basta

Scott Bowling

Janet Branchaw

Marguerite Brickman

Sara Brownell

Carly Busch

Alexa Clemmons

Liz Co

Jake Cooper

Erin Dolan

Miriam Ferzil

Brian Gibbens

Phil Gibson

Cara Gormally

Austin Heil

Angela Hodgson

Kelly Hogan

Mallory Jackson

Hannah Jordt

Laramie Denise Lemon

Stanley Lo

Tammy Long

Jenny McFarland

Jennifer Momson

Trish Moore

Cesar Nufio

Rebecca Orr

Megan Peterson

Deb Pires

Luanna Prevost

Carolyn Pucko

Kim Quillin

Michael Rodriguez

Heather Seitz

Justin Shaffer

Briana Timmerman

Ella Tour

Jacqueline Washington

We wish to thank the BioInteractive program, especially Melissa Csikari and Laura Bonetta, for hosting the early planning meetings, sharing the initial list of candidate LOs, introducing us to the education research scientists who advise the BioInteractive program at the Howard Hughes Medical Institute (HHMI), and for providing the BioInteractive Listserv.

Special thanks to Alexa Clemmons for sharing her expertise in survey design and knowledge of the Qualtrics Survey software.

Finally, thank you to the instructors listed below, as well as to those who wished to remain anonymous. Over 700 of you heeded the “call to action” and without you, this work would not have been possible.

Sarah Abboud

Sarah A. Arrington

Josh R. Auld

Jordan Baker

Michael Britton

Sue Ellen

DeChenne-Peters

Nancy Djerdjian

Eric Engstrom

Lawrence Hobbie

Justin Hoshaw

Hannah Jordt

Brian Kram

Kristin M. Lewis

Nicole Lynn McDaniels

Ryan J. Miller

Greg Podgorski

Kim Quillin

Margaret Saha

Cahleen Shrier

Rachel Smith

Drew Stenesen

Jessica D. Stephens

Alison Styring

Ann Thijs Charlie Willis

## Supporting information captions

S1 Data Set. All LOCS and HOCS candidate LOs with survey data and tags.

S2 Fig. Learning objectives sorted into eight bins identifies a “natural” 52.1% cut-off. NOTE: This was reviewed/endorsed by Groups 2 and 3.

S3 Table. Summary table of core learning objectives by unit for introductory biology for majors courses.

*Denotes a HOCS LO paired with two LOCS LOs or a LOCS LO paired with two HOCS LOs. S4 Text. Survey tool used to collect data on the essentiality of LOs.

S5 Fig. Unit learning objectives separated by LOCS and HOCS into low, medium, and high bins.

